# Breaking the cold chain: solutions for room temperature preservation of mosquitoes leading to high quality reference genomes

**DOI:** 10.1101/2025.07.03.662936

**Authors:** Fiona Teltscher, Petra Korlević, Edel Sheerin, Alex Makunin, Mara K. N. Lawniczak

**Affiliations:** Wellcome Sanger Institute, Hinxton, United Kingdom; James Cook University, Townsville, Australia

## Abstract

The Earth BioGenome Project (EBP) is a global endeavour to produce reference genomes for all described eukaryotic species. The majority of described species are arthropods, which tend to be small and require taxonomic expertise to identify to species level. Therefore, the ability to collect and preserve specimens in a suitable way for long read and Hi-C data generation using very simple approaches with minimal infrastructure is certain to be important in scaling up reference genome generation. Using *Anopheles* mosquitoes as an insect representative we evaluate how well different preservation liquids protect high molecular weight DNA, RNA, and nuclei for Hi-C when mosquitoes are held intact versus slightly squished. We find that squished samples stored in 100% ethanol and Allprotect held at room temperature for one week result in excellent preservation of both high molecular weight DNA and nuclei for Hi-C. Other tested buffers, including RNAlater, EDTA at several pHs, and DMSO Salt Solution (DESS) performed satisfactorily for long read data generation and RNA retrieval, but less ideally for Hi-C, which may have bigger negative impacts when aiming to generate data for organisms with larger genomes. Field collections requiring dry ice or dry shippers can be logistically challenging to arrange, are notoriously expensive, and DNA degrades rapidly if ultra-cold temperature is not maintained, which is devastating given how expensive and time consuming field work can be. Here we present multiple viable options for room temperature collection and/or shipment for arthropod samples. Further exploration across a broader range of species will hopefully enable cheaper and more widely available reference genome generation globally.

## Introduction

To date, only 1% of the earth’s described eukaryotic species have publicly available genome data and only 0.2% of species have high quality chromosomal reference genomes (https://goat.genomehubs.org/projects/EBP). Major advances over the recent years in long read sequencing quality from Pacific Biosciences (PacBio) and Oxford Nanopore Technologies (ONT), together with demonstrated scaleable extraction approaches across the tree of life (Howard et al. 2025) have highlighted the feasibility of the Earth BioGenome Project (Lewin et al. 2018, 2022) and projects like the Darwin Tree of Life (Darwin Tree of Life Project Consortium 2022), which has already released genomes for over 2000 species, are putting these advances into action. On the data generation side, challenges remain in scaleable assembly and curation, but perhaps the largest remaining challenge is securing sufficient funding. However, on the specimen collection side, there remain major challenges in identifying and collecting specimens at scale and collection challenges are made even more difficult by the need to collect and store with constant and reliable access to ultra cold temperatures. In practice, cold chain enabled specimen collection is managed either by collecting live specimens and returning with them to the lab where they can be rapidly processed, or through the use of dry shippers or dry ice in the field. However, all of these approaches add a logistical burden and considerable expense to field work, not to mention the risks should the cold chain method be compromised through dry ice evaporation or the dry shipper losing charge either during collection or shipping. Beyond these added expenses and risks, perhaps the greatest issue is that cold chain as a requirement precludes spontaneous collection when a species is encountered unexpectedly. Under current best practice guidelines for reference genome quality material (Lawniczak et al. 2025), when one encounters a specimen in the field but is either unable to keep it alive until getting to the lab or cold chain access is not available, the specimen should not be collected as it will not result in the kind of quality material needed for a reference genome. This guidance would be different if there were simple preservation buffers that protect HMW DNA, RNA, and nuclei from any specimen at room temperature and this would be a game changer for scaling up species collections as theoretically, any field work efforts could be “genome enabled”.

Previous work has set out to explore breaking the cold chain for vertebrate tissues (Dahn et al. 2022). This work explored ethanol, DESS, and DNAgard for several species and also Allprotect and RNAlater for fish. A variety of temperatures and tissues were explored, and not all preservatives were tested for compatibility with either long read sequencing or Hi-C. In short, most preservatives resulted in HMW DNA preservation but ethanol and DESS preserved higher molecular weight DNA better than DNAgard, which was also poor at preserving chromatin interactions for Hi-C sequencing. Vertebrate tissue is typically processed into smaller pieces at the time of sampling for various downstream approaches. The same may not be true for many arthropods, which are often small and easily collected as whole specimens with efforts to avoid morphological damage, as morphology may be required to reach an accurate species identification. Since both DESS and ethanol have long been known as good solutions for preserving DNA, we considered that these may not penetrate the insect cuticle rapidly enough to prevent DNA degradation and several years ago, began asking collaborators on our *Anopheles* Reference Genomes project (ENA Project ID PRJEB51690) to “lightly squish” their mosquito specimens in these solutions prior to storing and shipping at room temperature to best preserve DNA (Teltscher 2023). Here we more systematically explored compromising the cuticle versus leaving the specimen intact in several different preservation buffers and at two different temperature regimes, either storing the specimen in the freezer before subjecting it to a period of one week at room temperature, or simply storing it for one week at room temperature. In both cases, specimens were then removed from their preservation buffers and held at −70°C until further work.

The buffers we tested here include: 1) Allprotect Tissue Reagent (Qiagen), which is optimized to protect DNA in the 10-30 kbp range, RNA, and proteins; 2) DESS or DMSO Salt Solution (Seutin, White, and Boag 1991), which has been used across many species to protect DNA; 3) EDTA at three different pHs, because recent work exploring room temperature preservation of several aquatic species found that the primary protective ingredient in DESS was not the DMSO but the EDTA (Sharpe et al. 2020), and the same team showed that higher pH levels of EDTA can result in even greater HMW DNA recovery for some species (DeSanctis et al. 2023); 4) 100% ethanol, which is a common reagent used to preserve DNA at room temperature; and 5) RNA*later* (Thermo Fisher Scientific), which is a reagent that protects RNA in tissue including at room temperature. We tested each of these solutions on replicates of single *Anopheles* mosquitoes per collection tube held at room temperature for one week to ascertain which approaches result in high quality DNA, RNA, and nuclei preservation suitable for Hi-C.

## Materials and Methods

### Mosquito rearing and sample storage experiments

Seven to eight day old female *Anopheles coluzzii* N’Gousso strain laboratory reared mosquitoes fed on a 8% w/v fructose solution were collected with an electronic aspirator and anaesthetised using CO_2_. Batches of 6 to 12 females were transferred using forceps into a glass dish containing 100% ethanol, where they remained submerged for 1-3 min, followed by dabbing each individual on a Kimwipe to remove excess ethanol and transferring into 0.5 mL DNA LoBind tubes (Eppendorf). Each tube contained 400 µl of one of the following storage solutions: 100% molecular grade ethanol 200 proof (Fisher BioReagents), DESS (**D**MSO/**E**DTA/**S**alt **S**olution, comprising DMSO 20%, EDTA 0.25 M (Fisher BioReagents, crystalline powder dissolved in distilled water), NaCl to saturation, pH adjusted to 8 with HCl 5 mol/l or NaOH 5 mol/l from VWR Chemicals), 0.25 M EDTA pH 8, 0.25 M EDTA pH 9, and 0.25 M EDTA pH 10 (same 0.25 M EDTA solution as above with further NaOH pH adjustments), and Allprotect Tissue Reagent (Qiagen, abbreviated to Allprotect throughout). As Allprotect is extremely viscous, it was not possible to ensure exactly 400 µl was in each tube, but this was approximately correct. DESS was autoclaved and all EDTA solutions were filter sterilized for longer room temperature shelf life storage.

For each storage solution two treatments were prepared in sets of five replicates: “intact” in which the mosquito was picked up by a leg or the proboscis from the petri dish and placed into the tube without any physical disruption to the carcass, and “squished” in which a mosquito was similarly placed into the tube but also lightly pressed one time against the side of the tube using a plastic pestle, typically used for grinding tissues, in order to compromise the cuticle but leave the mosquito in one piece (Teltscher 2023). One set of replicates for each storage solution and treatment was stored at room temperature on a laboratory bench (20-25ºC) for one week until further processing. Samples receiving this treatment are referred to as “1wRT” (one week room temperature) throughout. A week was chosen to represent a reasonable amount of time to support collection in the field without cold chain and/or shipment to another destination without cold chain. A second set of replicates was first placed in a −20°C freezer for one week prior to being stored at room temperature for another week. Samples receiving this treatment are referred to as “ff-1wRT” (freezer first followed by one week room temperature) throughout. This design was intended to simulate a situation in which field collection had access to a −20°C freezer (e.g. live samples were brought back to a lab for processing) and to test if freezing made any substantive difference to initial preservation at room temperature. After one week at room temperature and prior to longer term storage, mosquitoes were removed from their storage solution with clean forceps (wiped with 70% ethanol in between), transferred into new 1.5 mL DNA LoBind tubes (Eppendorf), and stored in a −70°C freezer until DNA extraction. The removal of specimens from their storage liquid enabled streamlined extractions as no thawing of buffers was required and frozen specimens could enter directly into manual grinding for DNA or RNA extraction. Five snap frozen controls were also collected by submerging CO_2_ anaesthetised mosquitoes in 100% ethanol for 1-3 min, removing excess ethanol by dabbing each mosquito on a Kimwipe before placing them in 1.5 mL DNA LoBind tubes, and immediately storing in a −70°C freezer. All samples were stored at −70°C for at least one week before DNA extraction. The full experimental design is schematically represented in Figure 1A.

**Figure 1.**
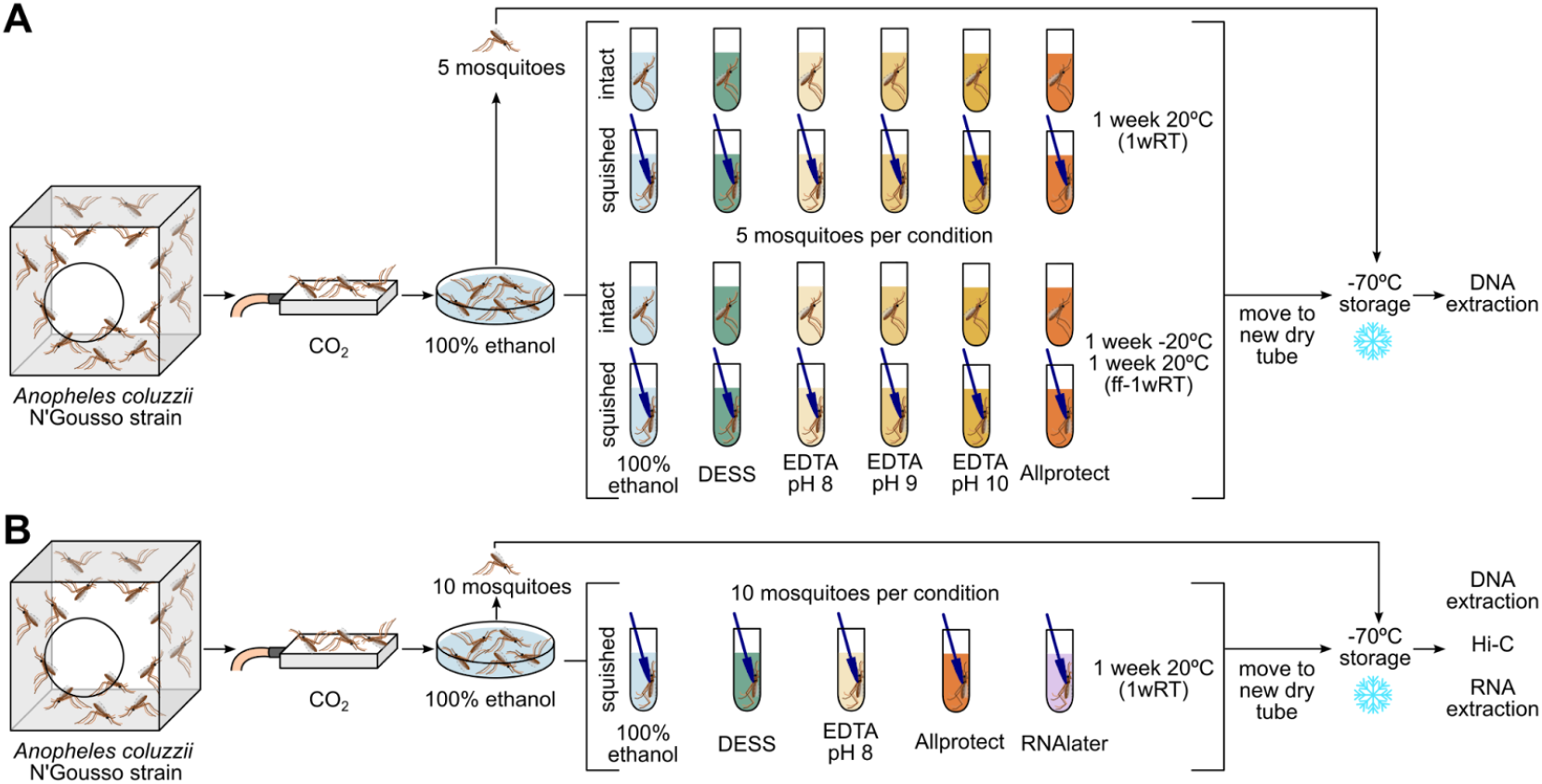
Schematic of our experiment simulating room temperature shipment of mosquitoes stored in various solutions. (A) Initial experiment on a variety of storage solutions and shipping conditions. These samples were extracted but only DNA quantification and fragment size assessment was performed. (B) Repeat experiment of a subset of storage solutions, for long read and Hi-C sequencing, and RNA quality assessments.

After assessing the best DNA preservation conditions, we selected the best performing subset of preservation liquids and also added in RNA*later* Stabilization Solution (ThermoFisher Scientific, abbreviated to RNAlater throughout) to test RNA preservation and chromatin configuration preservation for Hi-C sequencing, which are needed for new reference genome generation. One representative sample per condition was submitted for low input (LI) PacBio and Hi-C sequencing and a set of replicates for each preservation liquid was also evaluated for RNA quality. The selected storage solutions were 100% ethanol, DESS, EDTA pH 8, Allprotect, and RNAlater. For each storage solution 10 “squished” samples were prepared, five to be used for HMW DNA extraction, one for Hi-C, and four for RNA extraction. In addition, 10 snap frozen control mosquitoes were collected. We did not test the freezer first option in this second set of tests nor did we test intact mosquitoes given squished mosquitoes generally performed similarly or better. The experimental design is schematically represented in Figure 1B.

### DNA extraction and long read PacBio sequencing

DNA was extracted using the MagAttract HMW DNA Kit (Qiagen) on the KingFisher Apex Purification System (ThermoFisher Scientific). Briefly, samples are placed on dry ice, tissue is homogenized in lysis buffer using a plastic pestle and left at 25°C for 2 h to lyse, after which the volume is transferred to a KingFisher plate and the program performs two wash steps with buffers MW1, PE and nuclease free water, with a final 400 μl elution in buffer AE (Teltscher and Lawniczak 2025). Initial DNA quantification was performed using the Quant-iT PicoGreen dsDNA Assay (Invitrogen), followed by diluting each sample to about 0.25 ng/μl and re-quantifying using the Agilent Femto Pulse System using the 22 cm array and FP-1002-0275 genomic DNA 165 Kb kit following the “FP-1002-22 gDNA 165kb” method, which also assesses fragment lengths through smear analysis performed in the ProSize 4.0.0.3 data analysis software.

For the six extractions that were submitted for PacBio sequencing, the DNA was sheared to an average of 10-15 kbp using the Covaris g-TUBE system, where DNA is sheared by centrifugal force pulling it through a small orifice using a standard bench-top centrifuge. Following the Covaris manual, for each sample a total volume of 350 μl was centrifuge sheared. Due to the maximum volume of the Covaris g-TUBE being 150 μl, for each sample the volume was split into three centrifugation batches, 150+150+50 μl, and once each centrifugation was completed the batch was combined into the same 1.5 mL DNA LoBind tube. Each batch was centrifuged for a total of 6 minutes at 2,800 rpm, turning the g-TUBE every 1 min. Samples were concentrated and size selected with AMPure PB SPRI Beads (Pacific Biosciences) following the PacBio protocol “Preparing whole genome and metagenome libraries using SMRTbell prep kit 3.0” (102-166-600 April 2024) (PacBio 2025) with a few changes. The exact volume for each sample was roughly measured with pipette aspiration, then 3.1x 35% v/v beads were added and pipette mixed with a wide-bore tip. The sample-bead mixtures were incubated at room temperature on a tube rotor set to gentle rotation for 30 min, after which 1.5 ml of 80% v/v freshly prepared ethanol was used to wash the bead-bound DNA. DNA was eluted from the washed beads by adding 50 μl Buffer EB (Qiagen) and incubating in a benchtop shaker for 15 min at 37°C and 600 rpm.

Sheared and purified DNA was library prepped using the SMRTbell Prep Kit 3.0 (Pacific Biosciences, California, USA) as per the manufacturer’s instructions. PacBio libraries for all six samples were pooled together and sequenced on one Revio SMRT Cell. Sequencing was performed by the long read team in Scientific Operations at the Wellcome Sanger Institute.

### Hi-C sample preparation and sequencing

To evaluate whether these solutions are also suitable for preserving chromatin configuration, which is important for scaffolding reference genomes, a 1wRT squished specimen stored in each of 100% ethanol, DESS, EDTA pH 8, Allprotect, and RNAlater was submitted for Hi-C sequencing. Briefly, DNA was crosslinked and processed using the Arima-HiC v2 Kit (Arima Genomics) following the animal tissue protocol, after which Illumina libraries were sequenced on an Illumina NovaSeq X with 150 PE on the 25B flow cell aiming for >100X coverage per library.

### Genome assembly

The HiFi reads were first assembled using Hifiasm 0.19.8-r603 (Cheng et al. 2021) with the `--primary` option. Haplotypic duplications were identified and removed with purge_dups 1.2.5 (Guan et al. 2020). The Hi-C reads were mapped to the primary contigs using bwa-mem 0.7.17 (Li 2013). The contigs were scaffolded using the Hi-C data with YaHS 1.2.2 (Zhou, McCarthy, and Durbin 2023) using the `--break` option for handling potential mis-assemblies. The scaffolded assemblies were evaluated using PretextView 0.0.2 (*PretextView: OpenGL Powered Pretext Contact Map Viewer*).

### RNA extraction and quality control

DNA structure is known to be more stable than RNA, especially at room temperature, as its double stranded structure protects it from oxidation for longer, and thus we also assessed whether these preservation solutions were suitable for preserving RNA. Four 1wRT squished specimens stored in each of 100% ethanol, DESS, EDTA pH 8, Allprotect, and RNAlater were RNA extracted using the Dynabeads mRNA DIRECT Purification Kit (Thermo Fisher Scientific) following the protocol for solid plant or animal tissue on the KingFisher Apex. Each mosquito was ground with a pestle motor in 300 μl of lysis buffer and subsequently passed through a 21G needle 3-5 times. Homogenised samples were stored on dry ice until all samples were processed, and then thawed simultaneously on wet ice and quickly spun in an Eppendorf miniSpin benchtop centrifuge (1,200 rpm for 30 sec) before transferring them to the KingFisher Apex sample plate. Magnetic Dynabeads were cleaned by placing the required volume of beads in their provided storage buffer on a magnet, removing the supernatant, adding the same volume of fresh lysis buffer, and then transferring 50 μl of washed beads into a second plate on the KingFisher Apex. The beads were then collected by the magnet, added to the lysed sample and incubated for 10 min at medium shaking speed. DNA bound beads were then washed twice with 600 μl of wash buffer A, followed by two washes with 300 μl wash buffer B. RNA was eluted in 50 μl of 10 mM Tris-HCl pH 7.5 for 3 min at 70°C and immediately placed on ice. RNA was quantified and fragment sizes explored using the Qubit RNA High Sensitivity Assay Kit (Thermo Fisher Scientific) and the High Sensitivity RNA ScreenTape Assay for TapeStation Systems (Agilent). Routine DNase treatment was done using the TURBO DNA-free Kit (Invitrogen) to remove any remaining DNA. In short, 0.1 volume of 10X TURBO DNase Buffer and 1 μl of TURBO DNase Enzyme were added to the samples in a PCR plate, gently mixed on an Eppendorf ThermoMixer at 800 rpm, then incubated at 37°C for 20 min. The DNase was then inactivated by adding 0.1 volume of DNase Inactivation Reagent, briefly shaking at 800 rpm, and incubating for 5 min at room temperature before centrifuging the plates at 2,000 g for 5 min. The supernatant was transferred to a new plate and the RNA quantity was assessed by Qubit RNA High Sensitivity Assay Kit.

## Results

### DNA preservation across storage solution and conditions at room temperature

We initially evaluated six storage solutions (100% ethanol, DESS, EDTA pH 8, pH 9, or pH 10, and Allprotect), two sample handling conditions (intact, squished), and two storage approaches (1wRT, ff-1wRT) for high molecular weight (HMW) DNA preservation. DNA quantity and length estimates were assessed using a Femto Pulse instrument for every replicate and condition. Due to the large number of replicates across conditions, the extractions were completed over two days, so we only tested for statistically significant patterns in intact and squished replicates for the same storage solution as they were always extracted and purified the same day using the same reagents.

For the 1wRT setup, we compared estimated total DNA amounts in ng (noting there is natural variability in mosquito size and thus DNA yields), DNA amounts for fragments above 5 kbp length, and average DNA lengths in intact versus squished samples across all storage solutions (Fig. 2A). The 5 kbp cutoff was chosen because long read sequencing approaches can use a SPRI clean-up to eliminate molecules shorter than ∼5 kbp. While there seems to be a trend of slightly more DNA retrieved among squished samples in comparison to intact samples for most preservation solutions, the only significant difference in yield is found between intact and squished samples stored in DESS and EDTA pH 10, where squished mosquitoes have more DNA above 5 kbp. We also find that average DNA fragment size lengths are significantly greater among samples that are squished and preserved in DESS, EDTA pH 8, EDTA pH 9 and Allprotect in comparison to those held intact in the same solutions (Fig. 2B and Supplementary Table 1: Tab exp1).

**Figure 2.**
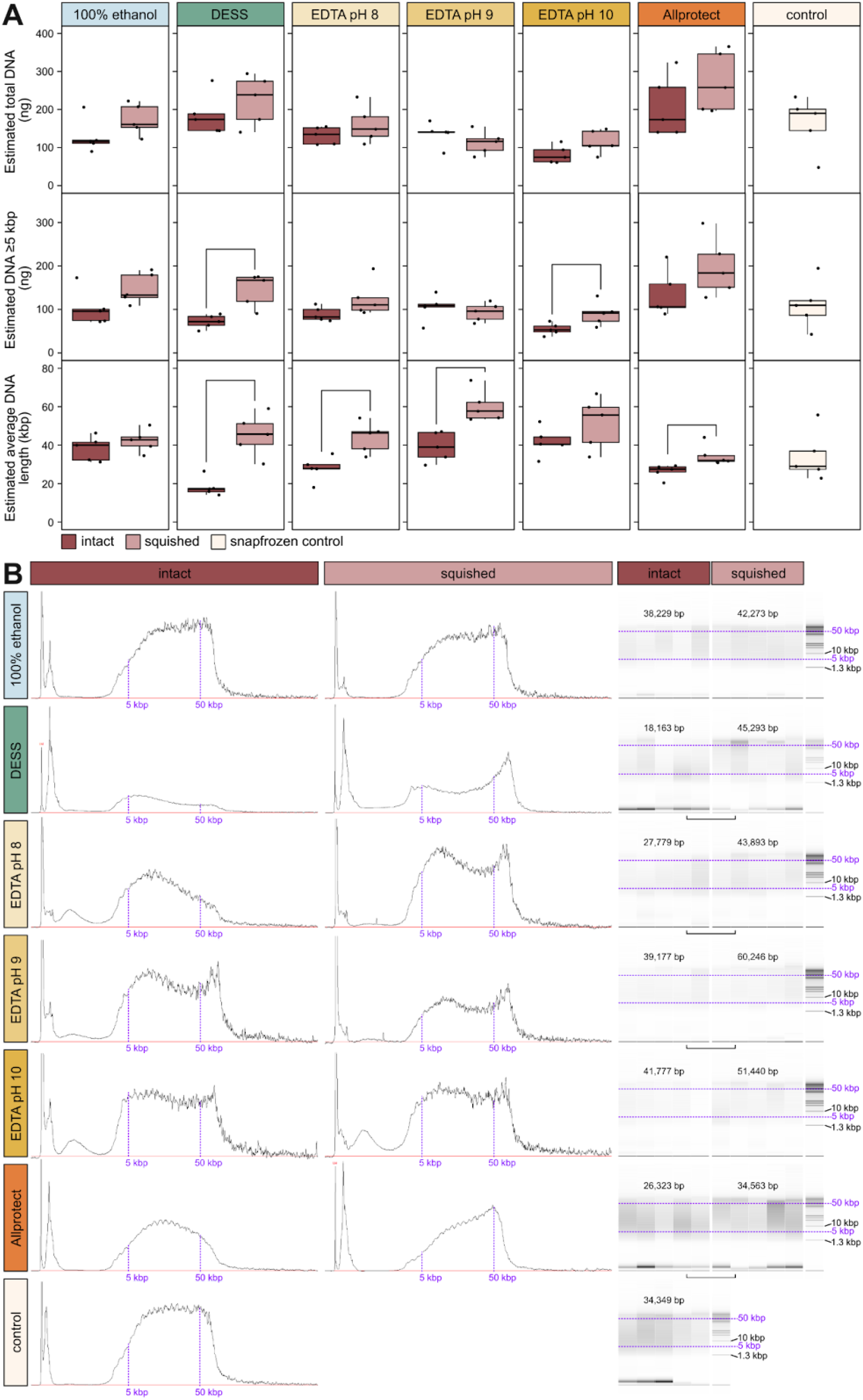
DNA quantities from intact versus squished samples held at room temperature for one week. **(A)** Estimated DNA yields (in nanograms) for 1wRT samples across all buffers for intact, squished, and snap frozen (control) replicates, top is total DNA, middle is only DNA above 5 kbp in length, bottom is estimated average DNA length (in kilobase pairs). Significant differences as assessed by t-test (p-value ≤0.05) are noted by brackets above (intact vs. squished). **(B)** Example Femto Pulse length profiles across buffers for a single intact (left) and squished (middle) replicate, as well as software approximations of gel smears for all replicates (right). Purple lines approximate the 5 kbp and 50 kbp lengths across samples. Average lengths across technical replicates are noted above the gel smears, with significant differences as assessed by unpaired t-tests represented by horizontal brackets. Heights are relative as each sample had a different dilution factor prior to loading (Supplementary Table 1).

Next we evaluated the ff-1wRT samples to understand if freezing the samples at −20°C in their storage solutions would promote solution penetration and subsequently help protect more and/or longer DNA (Fig. 3A). Like 1wRT, the overall trend in ff-1wRT samples also showed higher DNA yields in squished samples compared to intact samples, with DESS, EDTA pH 8, and Allprotect solutions resulting in significantly more DNA above 5 kbp. Average fragment lengths were also significantly larger in squished samples stored in DESS, EDTA pH 8, EDTA pH 9, and Allprotect than intact samples (Fig. 3B and Supplementary Table 1: Tab exp1).

**Figure 3.**
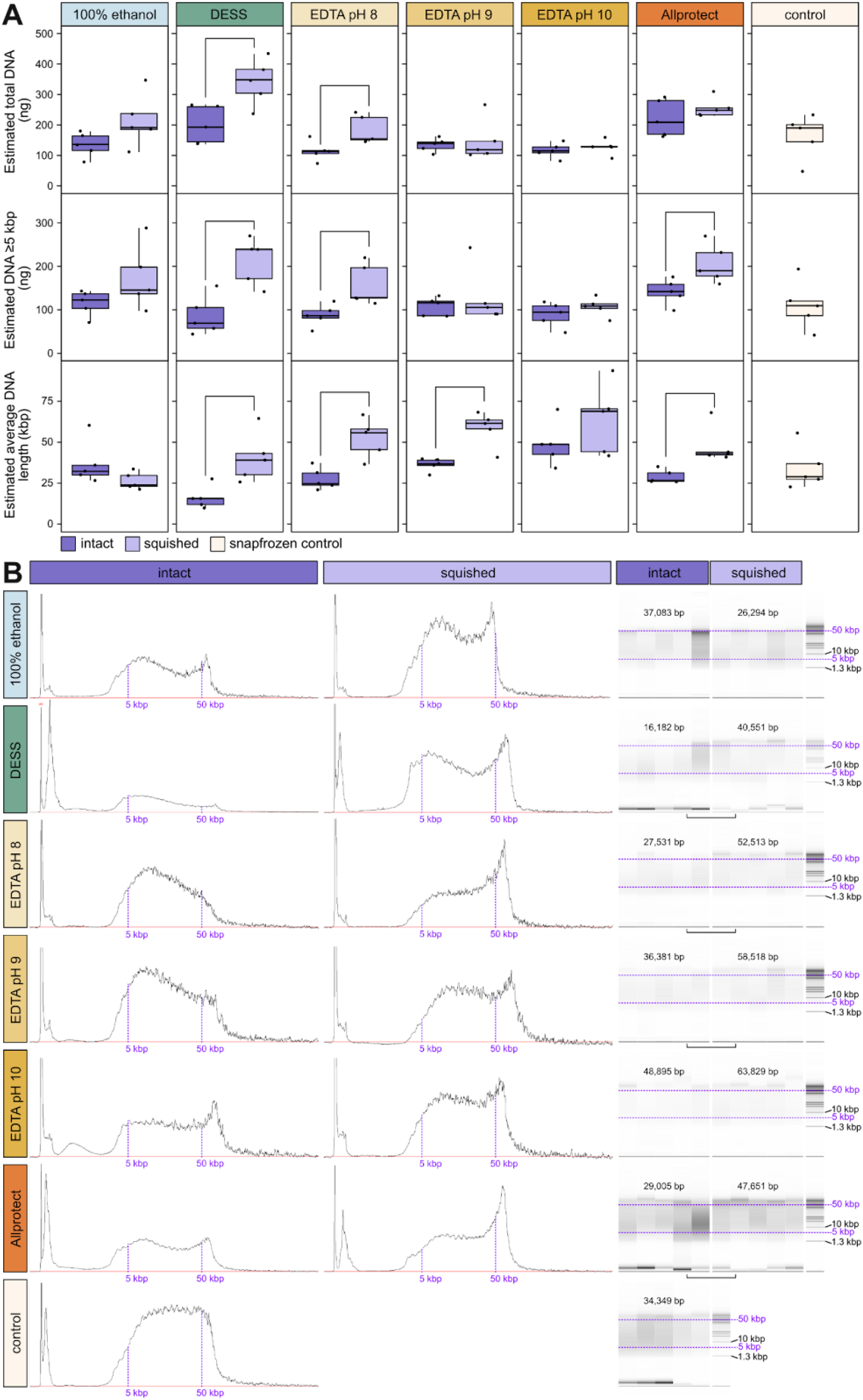
DNA quantities from intact versus squished samples that were first frozen then held at room temperature for one week. **(A)** Estimated DNA yields (ng) for ff-1wRT samples across all buffers for intact, squished, and snap frozen (control) replicates, top is total DNA and middle is only DNA above 5 kbp in length, bottom is estimated average DNA length (in kilobase pairs). Significant differences as assessed by t-test (p-value ≤0.05) are noted by brackets above (intact vs. squished). **(B)** Example Femto Pulse length profiles across buffers for a single intact (left) and squished (middle) replicate, as well as software approximations of gel smears for all replicates (right). Purple lines approximate the 5 kbp and 50 kbp lengths across samples. Average lengths across technical replicates are noted above the gel smears, with significant differences as assessed by unpaired t-tests represented by horizontal brackets. Heights are relative as each sample had a different dilution factor prior to loading (Supplementary Table 1).

Because the squished samples across storage temperatures showed a trend towards increased DNA yields (total and above 5 kbp), as well as longer retrieved fragments, we carried out additional t-tests comparing 1wRT and ff-1wRT squished replicates across solutions. We found that ff-1wRT DESS resulted in more total DNA than 1wRT, and 100% ethanol 1wRT resulted in longer DNA fragments than ff-1wRT (Supplementary Table 1: Tab exp1 bottom).

### Genome assembly outcomes across storage solutions

Current best practices in genome assembly consist of long read DNA sequencing together with Hi-C sequencing, ideally of the same specimen, to order and orient contigs. Given that all of our HMW DNA preservation conditions performed satisfactorily, with squished samples performing slightly better, we next prepared a test set to explore full genome assembly using the best performing subset of buffers including 100% ethanol, DESS, EDTA pH 8, Allprotect, and RNAlater. We selected only pH 8 among the EDTA options as it performed slightly better than the other two pHs and we also added in RNAlater for evaluation. We also selected only the 1wRT storage option because there were very minor differences between 1wRT and ff-1wRT, and the former does not require access to a −20°C freezer.

For this set of HMW DNA extractions, overall DNA yields were typically higher than for the previous set of extractions, within the normal range of variation that we observe, presumably due to differences in the size of our colony mosquitoes in any given generation. As this was a smaller sample set, we were able to extract everything in one day, and could thus compare DNA retrieval across all buffers at 1wRT and the snap frozen controls. We did not find any significant differences in the amount of DNA retrieved between the squished mosquitoes preserved at RT for one week in the solutions listed and the snap frozen controls, for total DNA and only DNA above 5kbp, which is highly promising for room temperature preservation of HMW DNA (Fig. 4A top, Supplementary Table 1: Tab exp2). Average fragment length was also comparable amongst experimental and control samples, with only 100% ethanol samples being significantly longer than the snap frozen control samples (Fig. 4A bottom). A single representative DNA extract for each preservation solution was chosen for shearing and sequencing (Fig. 4B). Although RNAlater and Allprotect showed more degraded profiles pre-shearing, they both sheared well to around a 10 kbp size along with all of the other preservation solution and the snap frozen control extractions (9,739 bp - 11,730 bp, Fig. 4B).

**Figure 4.**
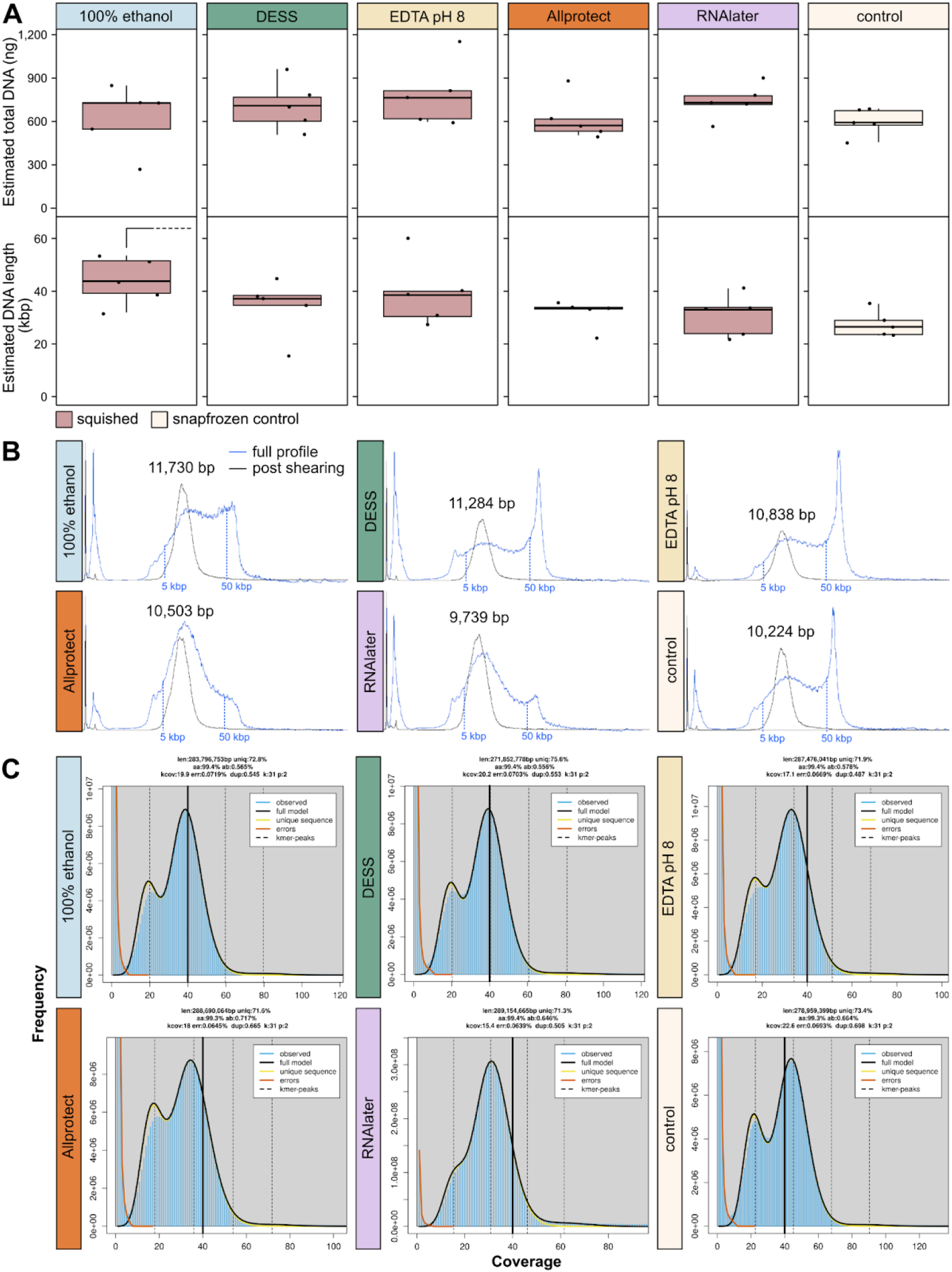
DNA quantities and assessments extracted from squished samples held at RT for one week. **(A)** Estimated total DNA (ng) for each of the preservation solutions and snap frozen controls (top) and estimated average DNA length (kbp) (bottom). Significant differences as assessed by t-test (p-value ≤0.05) are noted by half brackets above (storage solution vs. snap frozen control). **(B)** Femto Pulse traces for the samples selected for PacBio LI sequencing for each storage solution. The blue line is the original full profile for each sample and the black line is the same sample post g-TUBE shearing. The post-shearing peak size is noted above, with vertical blue dotted lines denoting the approximate location of 5 kbp and 50 kbp. **(C)** Resulting k-mer profiles for each sequenced PacBio library. Summary text above each plot summarizes haploid genome length (len), % unique sequences (uniq), percent homozygous k-mers (aa), percent heterozygous k-mers (ab), inferred coverage (kcov), read error rate (err), fraction duplicated reads (dup), k-mer size (k), ploidy (p). The colored lines showcase observed k-mer coverage frequencies (blue), the fitted model for all (black), unique (yellow), and error (red) k-mers. The thicker black line denotes a 40x coverage.

Each of the six sheared DNAs had PacBio libraries prepared and these were sequenced on a single Revio cell, aiming for > 15x coverage per haplotype per specimen. The *Anopheles coluzzii* genome size is approximately 250 Mbp, which led to this plexing decision. We achieved excellent sequencing results for each of the six libraries, with a typical N50 fragment length of > 10 kbp (Table 1) and GenomeScope profiles that suggested a successful sequencing run (Fig. 4C). This indicates that none of the preservation buffers caused DNA damage that would have resulted in reduced yields.

**Table 1.** PacBio sequencing and contig assembly summary for six samples sequenced using LI and one using ULI.

| Storage solution | k-cov | Haploid size | Repeat (%) | Het. (%) | yield | N50 | Contig length | Contig number | Contig N50 (L50) |
| --- | --- | --- | --- | --- | --- | --- | --- | --- | --- |
| 100% ethanol | 19.91 | 283,796,753 | 27.29 | 0.57 | 11,591,002,911 | 11,777 | 270,585,947 | 135 | 16,642,641 (5) |
| DESS | 20.20 | 271,852,778 | 24.48 | 0.56 | 11,261,944,002 | 10,891 | 271,115,124 | 174 | 15,936,013 (6) |
| EDTA pH 8 | 17.07 | 287,476,041 | 28.17 | 0.59 | 10,048,123,268 | 11,208 | 272,880,309 | 183 | 17,949,691 (6) |
| Allprotect | 17.95 | 288,690,064 | 28.47 | 0.73 | 10,610,020,181 | 10,452 | 273,620,267 | 165 | 14,159,000 (6) |
| RNAlater | 15.40 | 289,154,665 | 28.78 | 0.67 | 9,112,472,994 | 10,219 | 272,936,586 | 163 | 14,505,387 (6) |
| Control | 22.59 | 278,959,399 | 26.70 | 0.67 | 12,913,361,369 | 11,455 | 278,482,832 | 127 | 16,790,217 (6) |
| ULI | 19.49 | 248,552,781 | 21.21 | 0.74 | 10,109,927,058 | 8,187 | 262,372,469 | 846 | 1,003,577 (80) |

Due to the small size of the mosquitoes and the need to use an entire mosquito’s DNA extract for long read sequencing, for each preservation solution, Hi-C data was generated from a second individual. Therefore, we evaluated the performance of Hi-C data generated from squished specimens stored in each of the different preservation solutions using two approaches. First, we self scaffolded, meaning we used the long read data from the same preservation approach as was used for the Hi-C data (e.g. the two squished individuals that were both stored in 100% ethanol and used for long read and Hi-C sequencing). However, self scaffolding results were not particularly informative due to the high quality of the PacBio LI data for each specimen. Reconstruction of the genome for every preservation test resulted in 4-8 scaffolds in total but this was largely due to >1 Mbp contigs for >90% of the assembly from the long read data alone, so even a relatively low Hi-C signal was sufficient to merge these large contigs together (Table 2, Figure 5A).

**Table 2.** Hi-C scaffolding summary to newly sequenced PacBio LI samples and one previously sequenced PacBio ULI sample. For “self scaffolding”, the PacBio library and Hi-C library were from different individuals preserved using the same storage solutions. For the “ULI scaffolding” one ULI sample was scaffolded using Hi-C data from each of the different storage solutions.

| Storage solution | Hi-C coverage | Self scaffolding |  |  | ULI scaffolding |  |  |
| --- | --- | --- | --- | --- | --- | --- | --- |
|  |  | Scaffold number | Scaffold N80 (L80) | Scaffold N90 (L90) | Scaffold number | Scaffold N80 (L80) | Scaffold N90 (L90) |
| 100% ethanol | 504.1 | 112 | 2,927,161 (5) | 2,352,000 (6) | 316 | 2,556,046 (5) | 791,104 (15) |
| DESS | 377.5 | 150 | 4,135,150 (4) | 2,208,000 (5) | 561 | 3,625,167 (5) | 314,510 (41) |
| EDTA pH 8 | 367.0 | 155 | 19,127,350 (6) | 5,573,949 (8) | 694 | 4,251,698 (11) | 281,865 (45) |
| Allprotect | 447.0 | 140 | 3,228,000 (5) | 3,030,157 (6) | 348 | 2,168,088 (5) | 484,000 (20) |
| RNAlater | 502.8 | 136 | 3,358,947 (4) | 2,806,863 (5) | 441 | 2,411,669 (4) | 316,772 (37) |
| Control | 517.2 | 104 | 4,903,000 (5) | 4,514,151 (6) | 328 | 2,402,669 (5) | 645,855 (17) |

**Figure 5.**
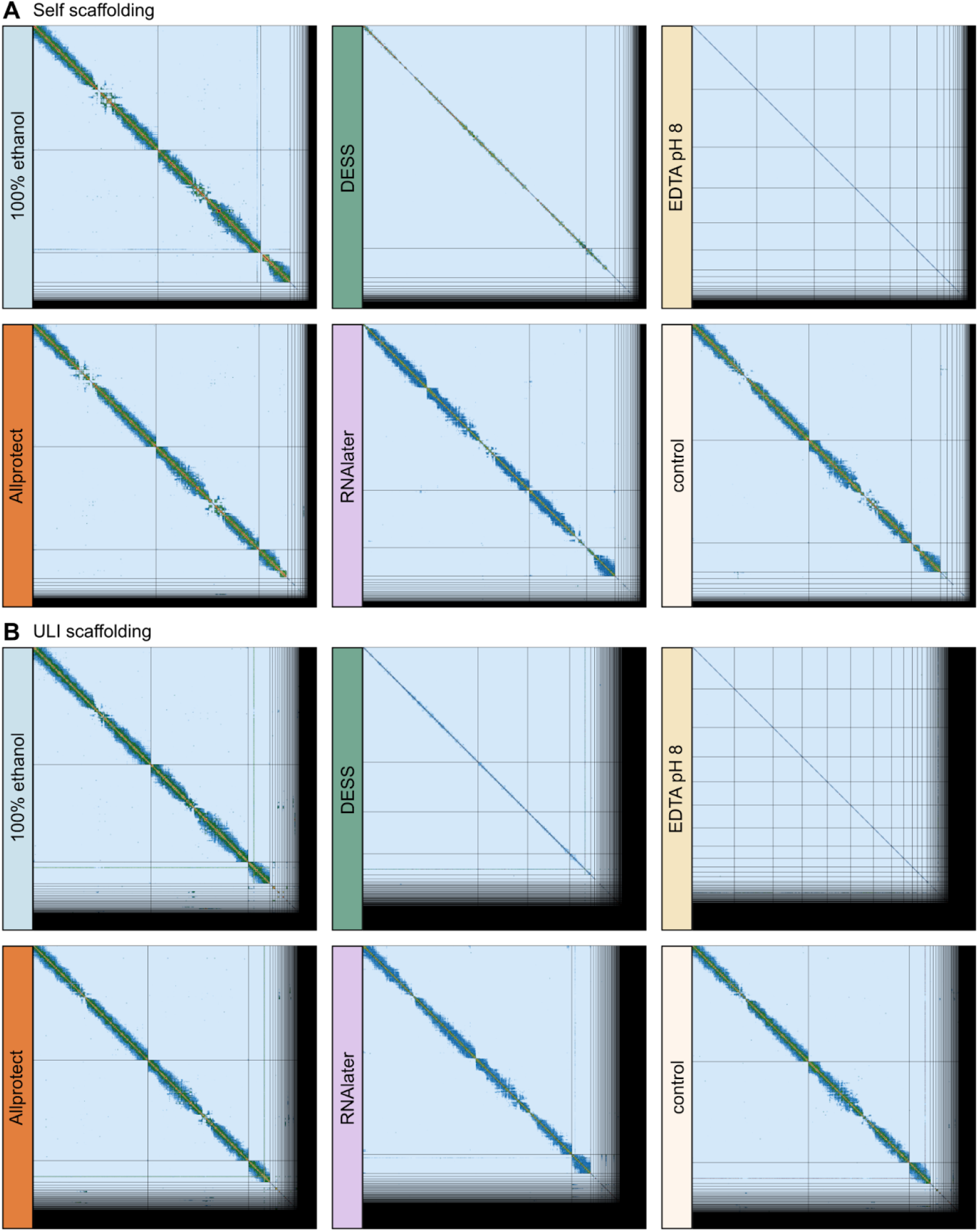
Scaffolded Hi-C plot for different storage solutions. (A) Self scaffolding. (B) ULI sample scaffolding. The grid for each preservation approach corresponds to the resulting scaffolds in decreasing size order, the color corresponds to the Hi-C contact frequencies - further from diagonal are more long-range contacts.

Second, we used each of the Hi-C datasets to scaffold a much lower contiguity long read assembly derived from a previously sequenced specimen from the same mosquito colony that was sequenced using PacBio Ultra Low Input (ULI) (Table 1 last row). PacBio ULI uses amplification to overcome low quantities of DNA, and has recently been replaced by PacBio AmpliFi. The ULI contig assembly has lower N50 fragment lengths (8,187 kbp) and the L50 (number of contigs in which 50% of the genome is covered) is at 80, whereas for all six LI samples, the L50 was 5 or 6. This much more fragmented ULI contig assembly provides a better assessment of the scaffolding success of each of the Hi-C libraries, as the contig assembly is in over 800 pieces as opposed to under 200, therefore the scaffolding outcome depends much more on long-range contact information from the Hi-C data. In this ULI scaffolding analysis, the most comparable conditions to the snap frozen control were 100% ethanol and Allprotect, with similar Scaffold N90 and L90 values (Table 2). DESS and EDTA pH 8 showed both decreasing long-range contact frequency (Fig. 5B) and decreasing efficiency in scaffolding the ULI sample when comparing the N80 (L80) and N90 (L90) scaffolds (in which we showcase how many scaffolds fit 80% and 90% of the genome, respectively, and note a larger jump in L90 for those preservation solutions) (Table 2). RNAlater was inconsistent, performing similarly to DESS and EDTA for scaffolding (Table 2), yet showing more long range contacts than these preservatives (Figure 5).

Finally, we also examined the quantity and quality of RNA retrieved from each of the preservation solutions. mRNA quantity was evaluated before and after DNase treatment, with a t-test comparing conditions to the snap frozen control finding that the average RNA amounts were significantly lower in 100% ethanol and Allprotect, but all preservatives resulted in similar average RNA fragment lengths (Supplementary Table 1: Tab exp2 bottom, Fig. 6). Evaluating RNA profile quality is difficult as insects do not reliably have the distinct rRNA bands that permit the use of the RNA Integrity Number (RIN), thus we inspected the profiles for visible signs of degradation. In comparison to snap frozen control, there are signs that DESS and EDTA pH 8 one week room temperature preserved samples show more degraded RNA, as does one ethanol preserved sample (Fig. 6B). In contrast, RNAlater and Allprotect, which are marketed as RNA preservation solutions by their respective providers, have RNA profiles similar to the snap-frozen controls (Fig. 6B).

**Figure 6.**
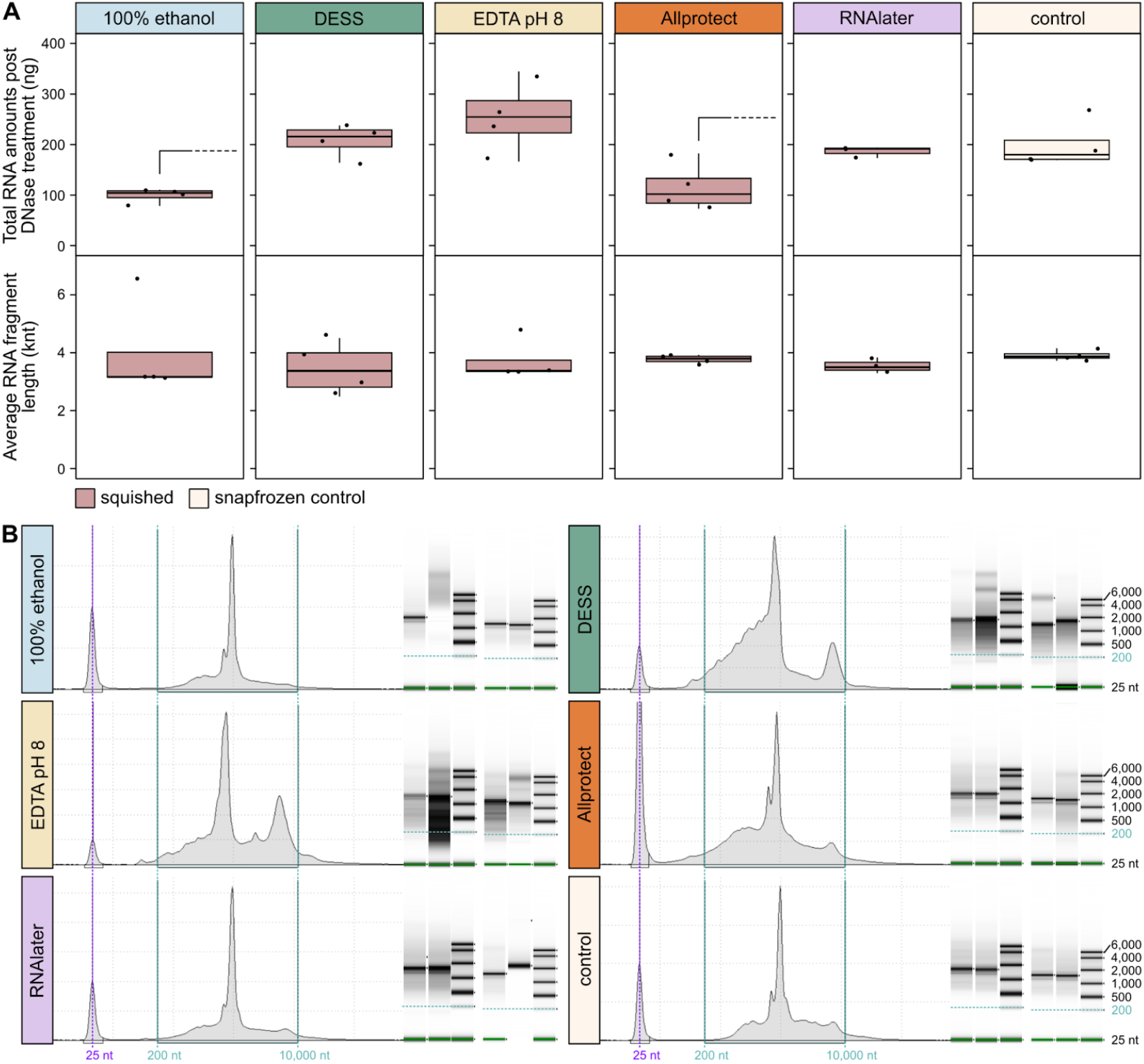
RNA retrieval from different storage solutions. (A) Four squished 1wRT samples per preservation solution with quantities measured with the qubit HS RNA kit after DNAse treatment (top) and average RNA length measured with the High Sensitivity RNA ScreenTape Assay for TapeStation Systems. Significant differences as assessed by t-test (p-value ≤0.05) are noted by half brackets above (storage solution vs. snafprozen control). (B) Example RNA profiles generated on the TapeStation. For each preservation solution, two replicates were run separately and gel representations of each of the four profiles together with their ladders can be seen to the right in each subplot. Purple vertical lines denote the 25 nt marker, with two vertical green lines denoting the 200 nt and 10,000 nt range, with the 200 nt line also being noted horizontally across the gel representations.

## Discussion

Snap freezing tissue and/or specimens and retaining them at ultra cold temperatures is considered best practice for long read, Hi-C, and RNAseq data generation, but snap freezing in the field is logistically challenging and expensive and requires preparatory work to ensure adequate access to dry shippers or dry ice. Even when collecting using ultra cold storage is possible, the material may need to be shipped elsewhere for data production, and shipping with dry ice or in dry shippers results in considerable expense as well as risk of cold chain loss. In our experience, shipping mosquito samples on dry ice from Cameroon to the UK would have been considerably more expensive (several thousand GBP) than the production costs of creating the reference genome for the species. As a result of cold-chain costs and risks, we have been trialing different room temperature preservation approaches to support cold chain free collection and shipping for insects. These trials led to the development of the squish method (Teltscher 2023) and also successful genome generation for the samples that were eventually shipped to the UK at room temperature from Cameroon (Nsango et al. 2023). However, we did not systematically compare different preservation approaches in a controlled fashion. Here, using colony mosquitoes reared in the UK, we evaluated a number of preservation solutions and storage conditions to more thoroughly explore what short-term cold chain free preservation approaches can lead to high quality reference genomes for insects.

We find that all preservation solutions and storage conditions tested here preserve sufficient HMW DNA for *Anopheles* mosquitoes held at room temperature for one week, and long read data generated from squished mosquitoes for each of the five different preservation solutions tested here was indistinguishable from long read data from a snap frozen control. The quantity of HMW DNA that can be extracted from a specimen is dependent on the size of the tissue/specimen, and the quantity of DNA that is needed is dependent on the genome size of the organism. Thus, while the specific preservation solutions used for these small mosquitoes did not matter for the quantity and quality of the resulting long read data, for larger genome organisms or smaller body sizes these small differences in quantity and quality may indeed matter. For most preservation solutions, when we explore molecules above 5 kbp, we see that squishing insects results in better preservation of longer molecules than storing insects intact, suggesting that squishing may enable more rapid penetration of all solutions at room temperature. More generally, we find that squishing mosquitoes in their preservation solution prior to room temperature storage for one week tends to preserve a bit more and longer HMW DNA. We also find that there does not seem to be a major impact of freezing tissues at −20°C prior to exposing them to a period of time at room temperature, but generally advise that best results are likely to be obtained by keeping specimens at the coldest possible temperatures at all times.

For a number of promising preservation solutions, we also evaluated the quality of Hi-C data for samples squished in these solutions. We find that squished samples in 100% ethanol and Allprotect performed comparably to the snap frozen control with respect to their ability to scaffold a relatively fragmented long read ULI contig assembly. The other buffers (RNAlater, DESS, EDTA pH 8) also achieved some scaffolding but generally performed less well. Of note is that we had >100x coverage of Hi-C data for each sample, if projects were to generate much less Hi-C data or begin with more fragmented contig assemblies or larger genomes, these latter buffers may perform much worse. In summary, RNAlater, DESS, and EDTA pH8 are likely not preserving insect chromatin conformation as well as 100% ethanol and Allprotect, and as a result, these latter two preservation buffers should be prioritised for room temperature storage if Hi-C is a necessary data type, especially if the target organism has a large genome.

RNA yields were similar across different preservation approaches, albeit slightly lower in 100% ethanol and Allprotect, and while quality is somewhat difficult to evaluate in insects, the buffers marketed for RNA preservation (Allprotect, RNAlater) were the most consistently similar to snap frozen controls. It is not widely appreciated but > 96% ethanol does preserve arthropod RNA although generally not as well as other solutions, especially for longer (> 1 week) periods at room temperature (Torres et al. 2019; Hasegawa, Techer, and Mikheyev 2021; Kono et al. 2016).

Given the results above, we recommend squishing arthropods lightly in either 100% ethanol or Allprotect to be the best “one size fits all” preservation buffers for short term (< 1 week) room temperature storage with a view to using such preserved specimens to create reference genomes. Unfortunately, Allprotect is expensive at approximately $10/mL (Qiagen website), and as it is extremely viscous, it is nearly impossible to dispense accurate volumes using the pump nozzle bottle it is supplied in. An alternative approach if cost is a consideration and if tissues are not in short supply is to split a specimen into two parts – one part preserved in 100% ethanol for DNA and Hi-C and one part (or some specific tissues of interest) into RNAlater for RNA sequencing, as RNAlater is about 10-fold cheaper than Allprotect. Additionally, if Hi-C is not needed, or the ambition is to sequence ultra long molecules, e.g. using Oxford Nanopore Technologies, then EDTA-based preservation approaches may be more appropriate as they tend to preserve longer molecules (Fig. 4). This requires further testing to ensure sequencing compatibility of EDTA preserved specimens with ONT. Other recent work evaluated the performance of ethanol and DESS as a tissue preservation buffer across ten species during thawing prior to HMW DNA extraction and found EDTA resulted in better protection of HMW DNA (Messner et al. 2025), however, Hi-C success was not evaluated in this work.

Benefits of squishing insects in their preservation solution is likely to extend to other arthropods outside Diptera, but further testing would be beneficial. For all preservation buffers, the tissue size-to-preservation buffer volume ratios are important – here we opted for 400 µl of buffer for single mosquitoes, which typically weigh about 3-5 mg. It is also important to note the tube size should be adequate to squish the tissues and fill the tube so the sample does not risk becoming stuck in the lid out of the preservation liquid during the shipping or storage periods. Here, we also removed specimens from their preservation liquid after the room temperature storage period and before longer term freezing at −70°C. In any case, people seeking to use preservation buffers may wish to consider whether removal from the preservation buffer is required or whether this can be done at the time of further work taking place. One advantage to holding tissues long term in their preservation liquid is if the long term storage freezer fails, the tissue is likely to still be protected.

In short, factors like tissue to buffer volume ratio, tissue type, and longer term ultra cold storage approaches should all be considered when preparing to collect and ship specimens of different species. Chordates and large invertebrates are typically dissected into different tissues and/or into small pieces to fit into storage tubes, and this may improve tissue penetration of preservation solutions, as well as make it possible to obtain HMW DNA, Hi-C and RNA from a single individual. Users should factor in their own circumstances to determine which approach is best for their particular needs. For example, if the species is relatively small and/or has a large genome, the amount of HMW DNA quantity that can be extracted might be near the boundary of what is needed for long read sequencing without amplification. Thus, even though they may not be statistically significant, trends towards small increases in yield observed here with certain buffers and conditions may be beneficial. Difficult to find species may result in only one specimen collected, in which case it may merit using Allprotect regardless of its cost. In other cases, for example if collecting very large numbers of specimens, preservation solution cost may be an important factor, in which case a combination of 100% ethanol and RNAlater might be the best option. To meet the grand ambitions of the Earth BioGenome Project, there is a major need to reduce costs associated with collecting and shipping specimens, including the high costs and inconveniences of cold chain collections and shipping. Here we present results supporting that short term room temperature storage in a wide range of buffers can lead to adequate preservation of insect material for reference genome creation. There may be merits in efforts explicitly collecting for reference genomes to use these preservation buffers even if cold chain access is available given the protection to the specimens these buffers afford together with the quality of the resulting data. Future work is needed to expand tests on different preservation approaches to other taxa and perhaps also to develop lower cost buffers that maximally future proof specimens not only for RNA, DNA, and nuclei but also for morphology, proteins, and metabolites, none of which were evaluated here but all of which are of interest.

## Acknowledgements

All authors and the work contained within were funded by Wellcome award 220540/Z/20/A to the Wellcome Sanger Institute. The Bill & Melinda Gates Foundation Award INV-009760 to MKNL to build high quality reference genomes from wild caught *Anopheles* mosquitoes was also instrumental in motivating the work. We thank Katharina von Wyschetzki for preliminary work to evaluate several preservation approaches discussed here. We thank Sanger Scientific Operations for completing all long read and Hi-C sequencing.

---

**Supplementary Table S1**. DNA and RNA QC values across all replicates. **Tab 1:** DNA quality control and relevant t-tests across conditions for six room temperature storage solutions (100% Ethanol, DESS, EDTA pH 8, EDTA pH 9, EDTA pH 10, Allprotect). **Tab 2:** DNA and RNA quality controls of five room temperature storage solutions (100% Ethanol, DESS, EDTA pH 8, Allprotect, RNAlater). In both tabs statistically significant p values are noted in italic and red.

